# ViR: a tool to account for intrasample variability in the detection of viral integrations

**DOI:** 10.1101/2020.06.16.155119

**Authors:** Elisa Pischedda, Cristina Crava, Martina Carlassara, Leila Gasmi, Mariangela Bonizzoni

## Abstract

Lateral gene transfer (LT) from viruses to eukaryotic cells is a well-recognized phenomenon. Somatic integrations of viruses have been linked to persistent viral infection and genotoxic effects, including various types of cancer. As a consequence, several bioinformatic tools have been developed to identify viral sequences integrated into the human genome. Viral sequences that integrate into germline cells can be transmitted vertically, be maintained in host genomes and be co-opted for host functions. Endogenous viral elements (EVEs) have long been known, but the extent of their widespread occurrence has only been recently appreciated. Modern genomic sequencing analyses showed that eukaryotic genomes may harbor hundreds of EVEs, which derive not only from DNA viruses and retroviruses, but also from nonretroviral RNA viruses and are mostly enriched in repetitive regions of the genome. Despite being increasingly recognized as important players in different biological processes such as regulation of expression and immunity, the study of EVEs in non-model organisms has rarely gone beyond their characterization from annotated reference genomes because of the lack of computational methods suited to solve signals for EVEs in repetitive DNA. To fill this gap, we developed ViR, a pipeline which ameliorates the detection of integration sites by solving the dispersion of reads in genome assemblies that are rich of repetitive DNA. Using paired-end whole genome sequencing (WGS) data and a user-built database of viral genomes, ViR selects the best candidate couples of reads supporting an integration site by solving the dispersion of reads resulting from intrasample variability. We benchmarked ViR to work with sequencing data from both single and pooled DNA samples and show its applicability using WGS data of a non-model organism, the arboviral vector *Aedes albopictus*. Viral integrations predicted by ViR were molecularly validated supporting the accuracy of ViR results. Additionally, ViR can be readily adopted to detect any LT event providing *ad hoc* non-host sequences to interrogate.

## INTRODUCTION

The transfer of genetic material between separate evolutionary lineages is a recognized event that occurs not only among prokaryotes, but also between viruses and eukaryotic cells (1). Somatic integrations of different viral species, among the best of known of which are the human papilloma virus, hepatitis B and C viruses and the Epstein-Barr virus, have been linked to genotoxic effects possibly progressing into cancer (2). Consequently, several pipelines have been developed to identify viral sequences integrated into the human genome using whole-genome sequencing (WGS) data (i.e. HIVID, SummonChimera, Vy-PER; HGT-ID, ViFi, VirTect, BS-virus-finder) (3–9). Each of these computational methods is differentially versatile in terms of data input format (i.e. RNA-seq, DNA-seq or bisulfite sequencing data) reference viral databases or customization opportunities, sensitivity and accuracy and CPU requirements, as previously reviewed (10). All of these pipelines have in common having been designed for the well-annotated human genome. Viruses can also integrate into the genome of germline cells. The persistence and the outcome of these integrations, which are vertically transmitted, depend on their effects on the fitness of the host. If deleterious, integrations are lost. If they occur in chromosomal locations that are not transcribed or lack regulative functions, integrations may persist and accumulate mutations. There are also examples of viral sequences being co-opted by the host and exerting beneficial functions (11). The existence of these Endogenous Viral Elements (EVEs) has long been known, with studies focusing mainly on EVEs from retroviruses in mammalian genomes (12, 13). The recent development of modern genomic sequencing approaches has opened to the study of non-model organisms. The genomes of organisms as different as arthropods (i.e. ticks, hematophagous and non-hematophagous insects), fish (i.e. zebrafish), snakes, birds, vertebrates (i.e. primates, mouse, rat, opossum) and plants were shown to host EVEs, which derive not only from DNA viruses and retroviruses, but also from nonretroviral RNA viruses (14–22). In these non-model organisms, EVEs range widely in numbers, for instance hundreds or less than 10 EVEs were identified in the genomes of *Aedes* or *Anophelinae* mosquitoes, respectively (20, 23). Additionally EVEs of non-model organisms tend to occur in repetitive DNA, mostly in association with transposable element (TE) sequences and in Piwi-interacting RNA (piRNA) clusters (20, 23, 24). These newly-identified EVEs have been increasingly recognized as important players in different biological processes such as antiviral immunity and regulation of expression (25, 26). However, rarely studies of EVEs in non-model organisms have gone beyond their characterization from annotated reference genomes. The lack of bioinformatic tools able to account for repetitive DNA when mapping WGS data to a reference genome is hindering our ability to detect integration sites different than those already annotated in the reference genome, thus to understand the widespread occurrence and polymorphism of EVEs in host genomes and testing hypothesis on EVE function. To ameliorate this issue, we have developed ViR, a new bioinformatics pipeline which improves the characterization of integration sites by solving the dispersion of reads in genome sequences that are rich of repetitive DNA. We benchmarked ViR with low- and high-coverage WGS data from *Aedes albopictus*, to date the mosquito species with the largest genome size and highest TE content among Culicinae (27, 28). We extensively evaluated ViR performance using WGS data from both single and pooled DNA samples, followed by molecular validation of predicted insertion sites. Importantly, ViR can be adopted to interrogate any genome for the presence of non-host sequences, which will facilitate studies of lateral gene transfer (LT).

## MATERIALS AND METHODS

### Pipeline implementation

The pipeline is composed of four scripts, which work in two modules. The first three scripts (ViR_RefineCandidates, ViR_SolveDispersion and Vir_AlignToGroup) work together to overcome the dispersion of reads due to intrasample variability (module 1). ViR_LTFinder is designed to run independently from the others, when testing for LT events of non-host sequences which have none or limited (defined by the user) sequence similarity to sequences of the host (module 2).

#### ViR_RefineCandidates

This script selects from a list of chimeric reads the best candidate pairs supporting a viral integration by filtering reads based on their sequence complexity, expressed as percentage of dinucleotides (default < 80%), imposing a minimum length recognized as viral (default 30 nucleotides) and removing mates that can align within a defined window in the reference genome (default 10000) based on Blast and BEDtools packages, respectively (29, 30) (Fig. 1A).

**Figure 1:**
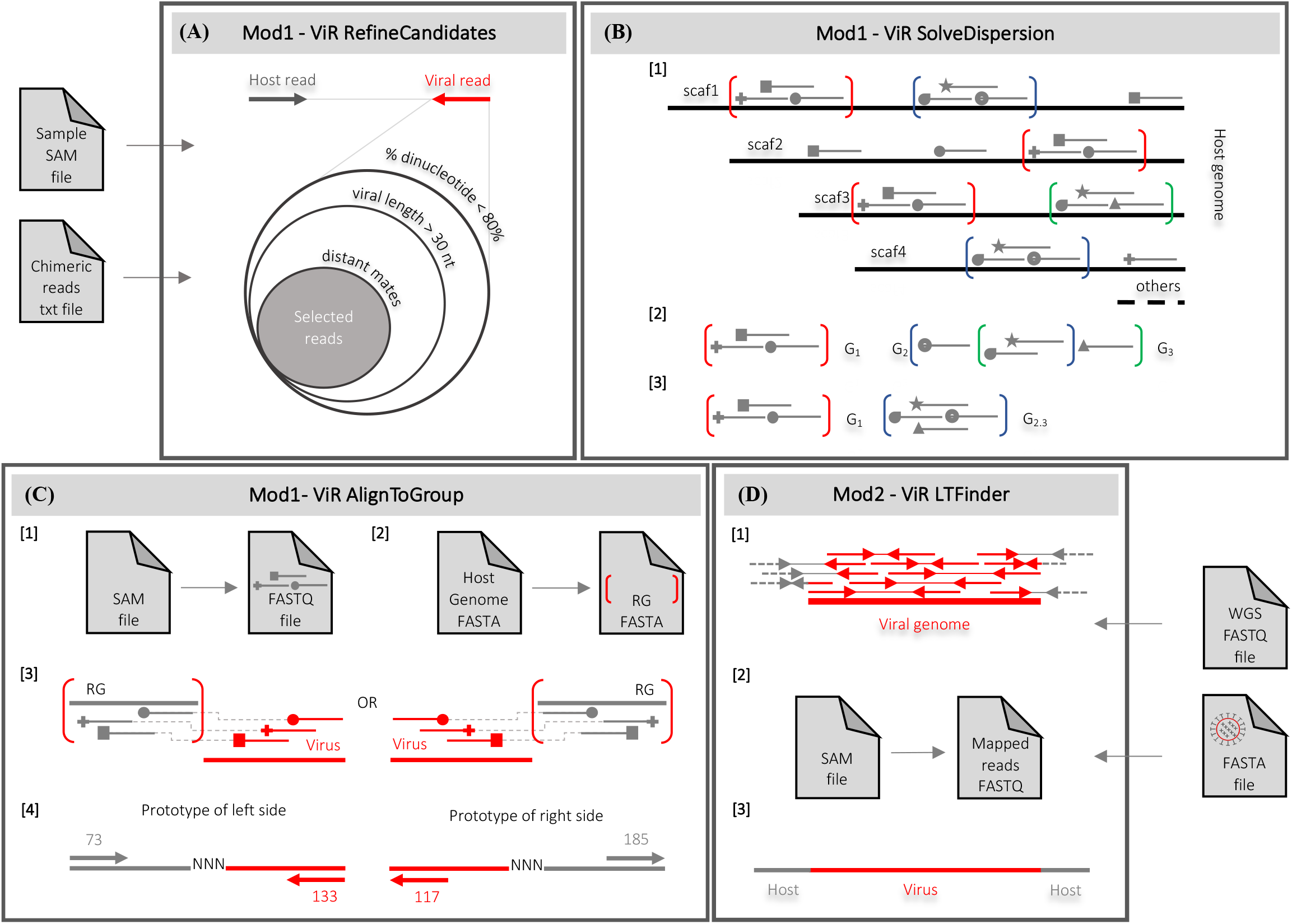
Overview of ViR. ViR comprises four main scripts, organized into two modules (Mod1 and Mod2): ViR_RefineCandidates, ViR_SolveDispersion, ViR_AlignToGroup and LT_Finder. **(A)** ViR_RefineCandidates filters the viral mate of the pairs of reads to identify the best candidate viral reads. Inputs for the script are the SAM file of the alignment of WGS data and a text file with the paired-end chimeric reads. Chimeric reads are pairs in which one read maps to the host genome (hereafter called host read, in gray) and one to a viral sequence (hereafter called viral read, in red) (10). Viral reads are identified using on a user-built database. Flags of alignment and sequence quality information are extracted for the identified best candidates. **(B)** ViR_SolveDispersion is designed to identify groups of host reads supporting an integration site in equivalent genomic regions (step 1). Equivalent means all genomic regions to which grouped reads can align. Read groups are compared in a pair-wise mode to merge groups sharing a certain percentage of reads (step 2). Remaining read groups support candidate viral integrations (step 3). **(C)** For each identified read group ViR_AlignToGroup extracts the reads from the SAM file of the selected reads by ViR_RefineCandidates (step 1) and the sequence of the equivalent region of the group in FASTA format (step 2). Then, reads are re-aligned to the sequence of the equivalent region and flags of alignment are used to identify the left and right side of the integration site (step 3). Examples of flags supporting the right (i.e. 73, 133) and left (i.e. 117, 185) ends of the integration site are shown (step 4). **(D)** ViR_LTFinder is used when searching for viral integrations from viruses for which no EVEs are annotated in the reference genome. WGS raw reads (WGS FASTQ file) are aligned to a viral genome(s) (FASTA file) (step 1). Aligned reads are extracted, converted into FASTQ (step 2) and used for local *de-novo* assembly (step 3). Aligned reads include mates in which both reads map to the viral genome (in red), chimeric reads (viral read in read and host read in dashed grey) and mates in which one is a soft clipped read (viral portion in red, the rest in dashed gray). Mates are indicated by continuous thin line.

Inputs for ViR_RefineCandidates are a text file of paired-end chimeric reads and the SAM file of the reads aligned to the host genome. Paired-end chimeric reads are pairs in which one read maps to the host genome (hereafter called host read) and its pair maps to a virus (hereafter called viral read) (10). The script is versatile and accepts as input the text file listing chimeric pairs, independently of the used tools. If no pair of reads pass the filtering steps, the following is printed on terminal: ‘No viral reads pass filters. Pipeline STOP!’. Otherwise, an output file is generated, which, for each pair of reads, collects information on both the viral and host read. Information include read flags, sequence quality data and relative positions of the two mates. Details of the content of the output file are listed in Supplementary Table 1.

#### ViR_SolveDispersion

This script solves the dispersion of host reads by grouping together reads that map to regions of the genome with the same sequence (Fig. 1B). Reads that are grouped are called “read groups”; regions of the genome to which these groups of reads can equivalently map are called “equivalent regions”. Equivalent regions are regions of the host genome with the same sequence because they contain the same repetitive element, or it is a sequence that has been erroneously assembled into different contigs or scaffolds. The script acquires as inputs a file listing all samples to analyze and the output directory of ViR_RefineCandidates. Reads mapping within equivalent regions are grouped together using the function “merge” from bedtools (Fig. 1B, step 1). Depending on the used blast settings, host reads may include also soft clipped reads (i.e. split reads in which a part of the read aligns to the host and the rest to a viral genome). The length of the equivalent region is defined by the user by setting the maximum distance among host reads and the minimum number of host reads within each region; defaults for these two parameters are 1000 base pairs (bp) and 2 reads, respectively. Then, identified read groups are compared in a pairwise mode in an iterative process in which read groups sharing more than a user-defined percentage of reads are collapsed in one; default percentage of reads shared between two read groups to be merged into one has been set to 80% (Fig. 1B, step 2). This procedure allows to identify the best candidate anchor genomic region of a candidate viral integration site (Fig. 1B, step 3). Each read group supports a candidate viral integration. Within each candidate, all viral reads correspond to the same viral species. Options to describe the genomic context of each region are available by proving adequate input files, such as the fasta file of the repeats of the host genome and/or the bed files of EVEs and piRNA clusters annotated in the reference genome.

#### ViR_AlignToGroup

This script predicts the right and left sides of the integration site by realigning reads supporting each candidate viral integration against the sequence of the equivalent region. First, for each candidate, this script extracts the host reads with their viral pair from the SAM file of the reads analyzed by ViR_RefineCandidates using the command-line utility “grep” (https://www.gnu.org/software/grep/manual/grep.html); the SAM file is converted into a BAM file using the function “view” of SAMtools (31); the BAM file is converted into a FASTQ format using the function “bamtofastq” from BEDtools (29) (Fig. 1C, step 1). Then, the script obtains the sequence of the equivalent region in fasta using the BEDtools function “getfasta” (29) (Fig. 1C, step 2). Reads from step 1 are re-aligned to the sequence of the equivalent region using “bwa mem” with default parameters (32). By taking advantage of the flags of alignment of each read of all chimeric pairs and eventual soft clipped reads, the left and right sides of the integration point can be predicted using Trinity (Fig. 1C, step 3). Even if no assemblies are created, flags of alignment are used to predict the direction of the integration sites (https://broadinstitute.github.io/picard/explain-flags.html) (Fig. 1C, step 4).

#### ViR_LTFinder

This script is designed to test for an integration from non-host sequences which have no, or a user-defined percentage of, similarity to host sequences. WGS reads are mapped to a selected non-host sequence (i.e. an entire genome or selected portions) using “bwa mem” with default parameters (32). Mapped reads are extracted using the function “view” of SAMtools (31) (Fig. 1D, step 1). These reads include pairs in which both mates map to the non-host sequence, chimeric reads and soft-clipped reads. The aligned reads are converted into FASTQ format using the function “bamtofastq” from BEDtools (29) (Fig. 1D, step 2) and used for *de-novo* local assembly using Trinity (33) (Fig. 1D, step 3). A consensus sequence is built if any instances of LT are identified. Output of ViR_LTFinder include files for visualization of the aligned reads using the Integrated Genomics Viewer (IGV) tool (34).

### Estimating the gain in solving read dispersion

The utility of ViR in solving read dispersion was assessed using the concept of the ‘Gain Index’ parameter from the Information Theory (35). This index estimates the weight that each attribute has in building a classification of entities, given n entities each defined through various attributes (35). The maximization of the Gain index reflects the power of an attribute in segregating entities to different classes. In our case, we used this index to evaluate the gain of enclosing in a single ‘equivalent region’, reads that had been originally assigned to different loci, our classes. The attribute is the “equivalent region’ identified by ViR_SolveDispersion and the entities are all the reads supporting candidate integrations (i.e. output of ViR_RefineCandidates).

The gain of running ViR is estimated through the formula:

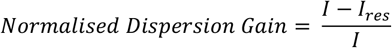

Where *I* is the entropy.

Entropy is the dispersion of the reads supporting each locus ID in a sample and it is quantified by the formula:

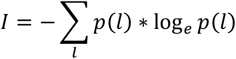

where *I* is the locus ID, meaning the host genomic coordinates of the candidate integration identified before ViR. *p*(*l*) is the relative frequency of the reads assigned to the locus ID *l*. Entropy is 0 when only one locus ID (i.e. one candidate integration site) is identified in the sample (i.e. WGS dataset). Entropy is > 0, when more than one locus ID is identified in the sample.

And *I_res_* is the residual information.

The residual information is defined by the formula:

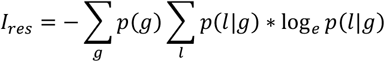

Where *g* is the ID of the equivalent region identified by ViR_SolveDispersion, *p*(*g*) is the relative frequency of the reads in the equivalent region *g* and *p*(*l*|*g*) is the relative frequency of reads assigned to the locus ID *l* in the equivalent region *g*.

The difference between I and *I_res_* is evaluated for each sample. The ratio of the difference with respect to the initial entropy is calculated to normalize across samples, which were obtained using different experimental set ups (i.e. WGS from single or pools). This operation results in a *Normalised Dispersion Gain*, which ranges between 0 and 1. The closer the value of *Normalised Dispersion Gain* is to 0, the higher is the gain of ViR in solving the dispersion of reads. To favor intuitive interpretation of results, we show Solve Dispersion Gain as:

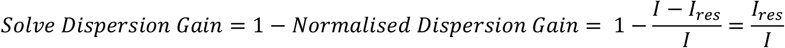

Values of *Solve Dispersion Gain* > 0 are found when ViR was able to identify a unique equivalent region for at least two different reads previously assigned to two different loci ID. The higher the value of *Solve Dispersion Gain*, the higher is the performance of ViR.

### Molecular validation of ViR-predicted integration sites

We run ViR on WGS data of *Ae. albopictus* and identified a total of seven candidate viral integrations absent in the reference genome assemblies, AloF1 and AalbF2 (27, 28). PCR primers were designed on the basis of these candidates (Table 1) and used on DNA extracted from different mosquitoes than those used as source of WGS data. *Aedes albopictus* mosquitoes are reared at the insectary of the University of Pavia as previously described (23). Genomic DNA was extracted from individual mosquitoes using the Promega Wizard^®^ Genomic DNA Purification Kit, according to manufacturer’s protocol. PCR reactions were carried out with the DreamTaq Green PCR Master Mix (ThermoFisher) using 1 μl of genomic DNA. Amplified bands were purified with ExoSAP-IT kit (ThermoFisher) and send to Macrogen (Madrid, Spain) for Sanger sequencing. Sequences were analyzed with Bioedit (36).

**Table 1.**
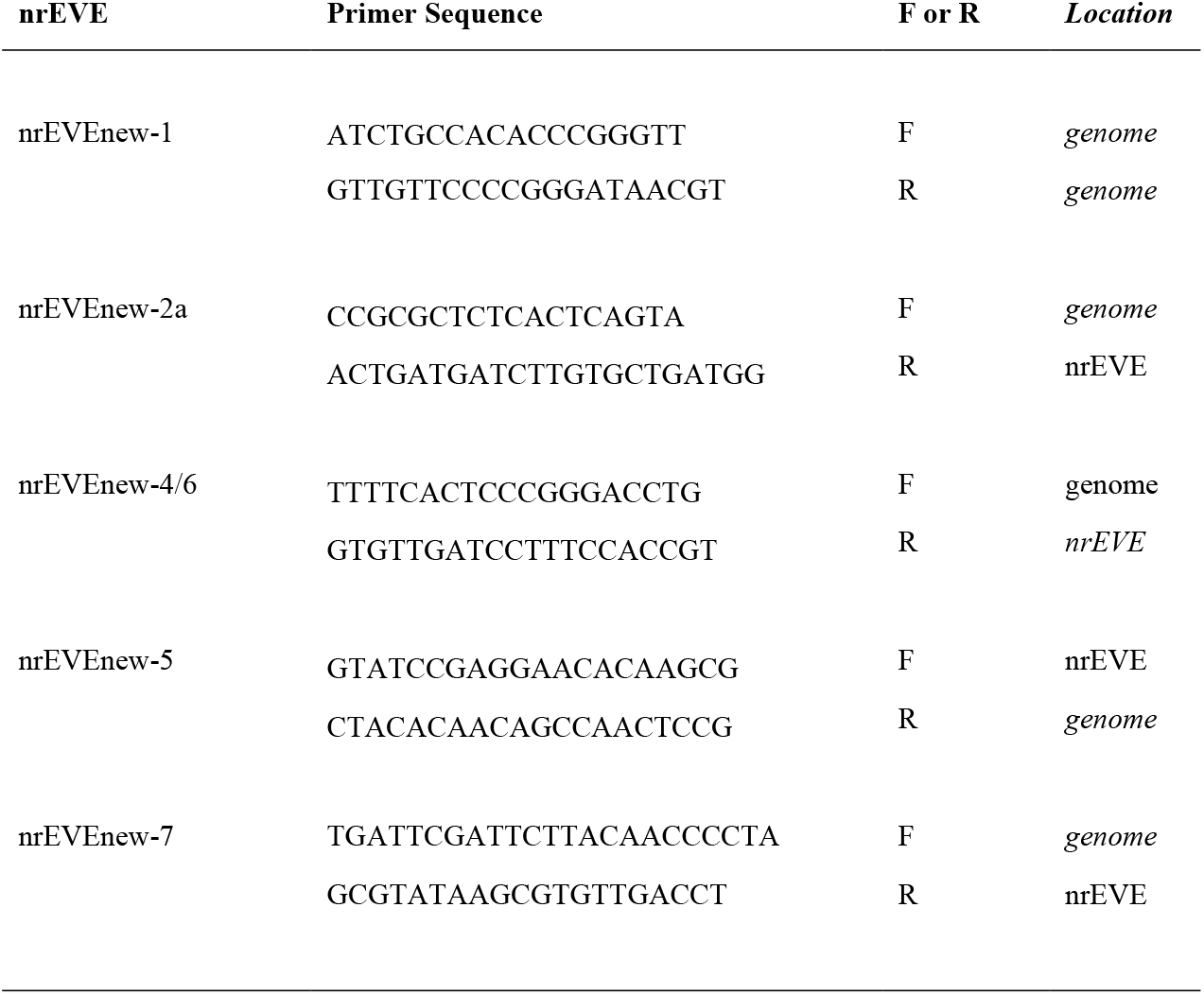
List of PCR primers used to confirm ViR-predicted viral integrations from WGS data of *Aedes albopictus*.

## RESULTS

### Overview of the ViR pipeline

ViR is designed to solve the dispersion of reads due to an integration site occurring in repetitive or misassembled regions of a genome. Any of the currently availed tools to identify viral integrations from paired-end sequencing data operates by selecting chimeric pairs, in which one read maps to the host genome (the host read) and the other to the viral genome (the viral read) (10). If the integration site occurs in a repeated region or in a genomic region that has been assembled into different contigs or scaffolds, host reads may be distributed across these “equivalent” genomic positions, and, as a consequence, the signal for the integration site, expressed in terms of host reads coverage, may not reach threshold of detection. To ameliorate this, we developed ViR, a pipeline composed of four scripts divided into two modules (Fig. 1).

Using the script ViR_SolveDispersion, reads that map to equivalent regions of the genome are grouped together (Fig. 1B). These groups represent anchor genomic regions where candidate viral integrations occurred. Additional information on these regions can be collected by providing input files with custom data (i.e. mapping coordinates of EVEs annotated in the reference genome, of piRNA clusters, of TEs, etc.). This additional information is useful to distinguish between polymorphisms of EVEs *versus* new integrations from the same virus. The best candidate pair of reads supporting a viral integration are identified by running the script ViR_RefineCandidates (Fig. 1A), which parsed viral reads through different filtering steps. This procedure should be performed before running the script ViR_SolveDispersion to reduce the computational time to solve the dispersion of host reads. Following ViR_SolveDispersion, the sequence of each equivalent genomic region is extracted, and host reads are re-aligned to it. This procedure allows to predict the left and right sides of the integration site through the use of alignment flags (Fig. 1C).

ViR_RefineCandidates, ViR_SolveDispersion and Vir_AlignToGroup work together to ameliorate the detection of integration sites by solving the dispersion of reads due to intrasample variability (module 1).

The pipeline comprises an additional script, ViR_LTFinder, which runs independently from the others (module 2). This script uses WGS data to map against a non-host sequence (i.e. entire genomes, genes, transposable elements). Aligned reads are extracted and used for *de novo* local alignenment using Trinity building a consensus sequence for the LT event (Fig. 1D). Eventual similarities between the tested non-host sequence and sequences of the host may lead to unbiguities, for instance if searching for integrations from a viral species for which EVEs are already annotated in the host genome, thus users have to decide a threshold for the percentage of similarity between their non-host sequence and eventual host sequences to avoid missinterpretation of results from ViR_LTFinder.

### Benchmarking ViR

ViR was run using WGS data from the arboviral vector *Ae. albopictus*. *Aedes albopictus* has the largest mosquito genome sequenced to date, the majority of which is repetitive DNA (27, 28). For *Ae. albopictus*, three genome assemblies are available, which differ greatly in the number of scaffolds and are all larger than the expected size of the genome based on cytofluorimetric estimates, suggesting duplications (27, 28, 37). We selected to run ViR using two assemblies, AaloF1 and AalbF2, which have 154782 and 2197 scaffolds, respectively. Hundreds of EVEs from various taxonomic viral categories are annotated in AaloF1 and AalbF2 (23, 28), giving further complexity.

We generated WGS data from DNA of both single and pool DNA samples and at different coverage to compare the accuracy of ViR across different conditions (Fig. 2A). Single samples consisted in twenty-two mosquitoes whose genomes were analyzed independently. WGS libraries were generated and sequenced on an Illumina HiSeqX platform as previously described (38). Single samples were sequenced at an approximate coverage of 30x. Pool samples were 6 samples consisting each of the DNA from 30 mosquitoes. Three samples were sequenced at an approximate coverage of 30x and called pool30; the remaining samples were sequenced at an approximate coverage of 60x and called pool60. A list of chimeric pairs, indicative of candidate integration sites, were obtained for each sample running Vy-PER (3) with a custom-made database of viral genomes (Supplementary Table 1). Using 4 CPUs and16 Gb RAM per sample, VyPER produced results in between 5-33 hours in AaloF1 and 6-68 hours in AalbF2 (Table 2). Chimeric reads identified by VyPER are spread across different regions of the host genomes (Fig. 2A) and viral reads include homopolymers or low-complexity sequences.

**Figure 2:**
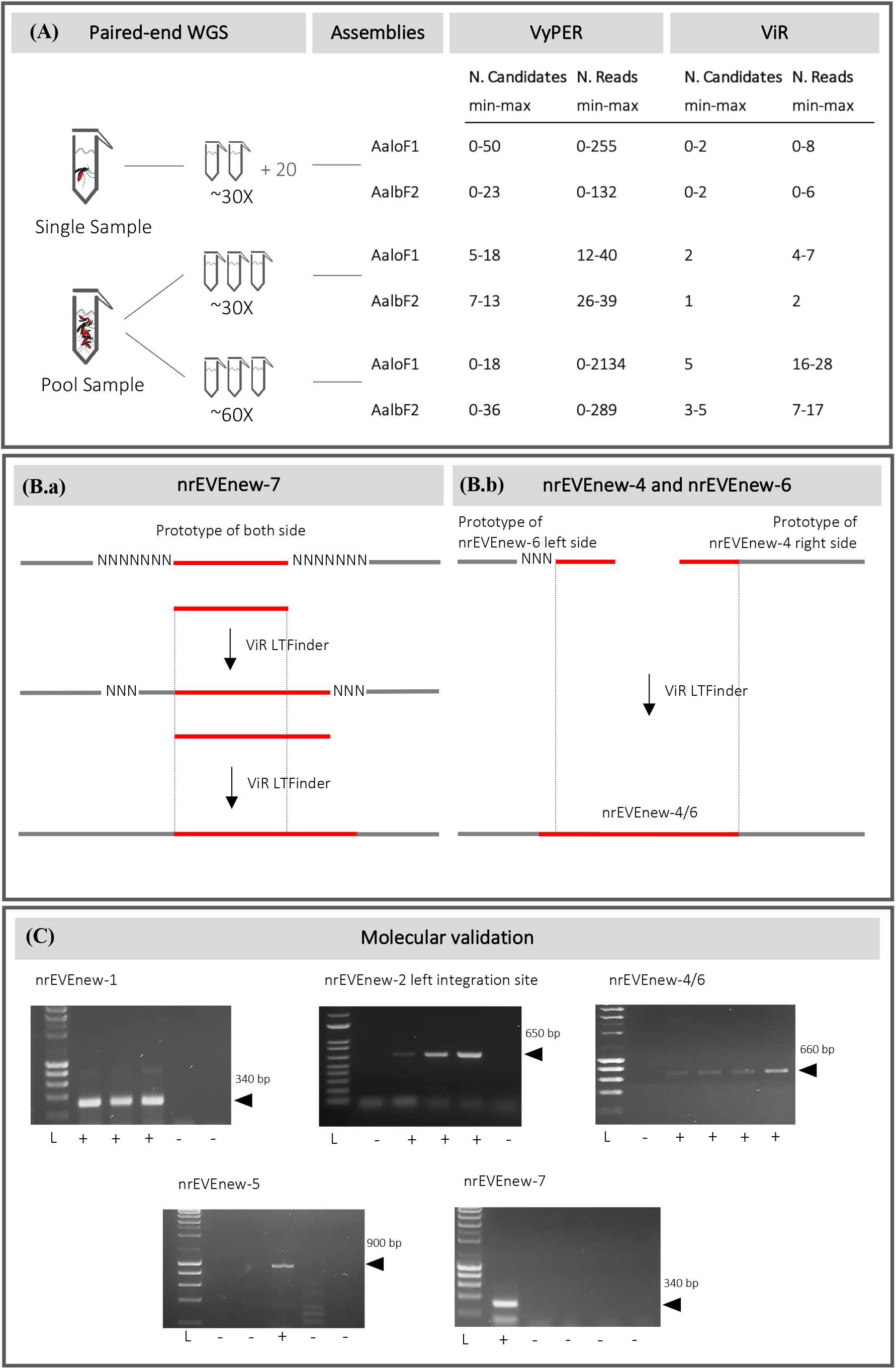
Benchmarking ViR. **(A)** Whole Genome Sequencing (WGS) data were generated from DNA of both single and pool DNA samples and at different coverage to compare the accuracy of ViR across different conditions. We selected to run ViR using two assemblies, AaloF1 and AalbF2 (28, 29) with a custom database of viral genomes (Supplementary Table 2). A list of chimeric pairs, indicative of candidate integration sites, were obtained for each sample running Vy-PER (3). Module 1 of the ViR pipeline was run using the chimeric reads identified by Vy-PER to solve read dispersion, resulting in a total of seven candidate viral integrations across all samples. **(B) (a)** ViR_LTFinder was run with WGS data and the viral sequence of nrEVEnew-7 generating an extended consensus (step 1). ViR_LTFinder was run a second time using this elongated viral sequence as input (step 2), which led to the identification of both integration sites (step 3); **(b)** ViR_LTFinder was run with WGS data and the viral sequences of nrEVEnew-4 and nrEVEnew-6 showing that they correspond respectively to the right and left regions of the same integration. **(C)** Molecular validation of the five candidate viral integrations. Each lane is the result of a PCR on mosquito genomic DNA with primers that were design to check the left integration site for nrEVE-new-2, and to check both the left and right integration points for the others viral integrations. ‘+’ indicates the presence of the viral integration and ‘-’ the absence.

**Table 2.** Per sample computational and time cost for VyPER implementation in different experimental conditions.

|  | Single sample |  | Pool30 |  | Pool60 |  |
| --- | --- | --- | --- | --- | --- | --- |
|  | AaloF1 | AalbF2 | AaloF1 | AalbF2 | AaloF1 | AalbF2 |
| CPU | 4 | 4 | 4 | 4 | 4 | 4 |
| Memory | 16 Gb | 16 Gb | 16 Gb | 16 Gb | 16 Gb | 16 Gb |
| Mean Time | 8 h | 11 h | 5 h | 6 h | 33 h | 68 h |

#### Testing module 1

ViR_RefineCandidates and ViR_SolveDispersion were run for all samples using both AaloF1 and AalbF2 (Fig. 2A). Parameters set in ViR_RefineCandidates included filtering out viral reads with homopolymers (any combination of dinucleotides higher than 80%), viral matches shorter than 30 nucleotides and distance between read pairs higher than 10000 nucleotides, considering the maximum DNA fragment length produced in our WGS data. Parameters set in ViR_SolveDispersion included defining a read group when at least two host reads aligned within a window of 1000 nucleotides and merging read groups than shared more than the 80% of reads. Publicly available annotations of EVEs, piRNA clusters and transposable elements were used to define the genomic context of each equivalent region (23, 28).

When ViR was run with AaloF1, five and two candidate viral integrations were identified in pool60 and pool30 samples, respectively, with a maximum number of reads per sample of 28. Viral integrations were also identified in eight out of the 22 tested single samples, with a maximum number of reads per sample of eight (Fig. 2A). When ViR was run with AalbF2, we identified a maximum of five viral integrations in pool60 samples and one in pool30 samples with a maximum number of reads per sample of 17. As for AaloF1, eight of the 22 single samples showed candidate viral integrations, with a maximum number of reads per sample of six (Fig. 2A).

Overall across all samples, a total of seven integrations, all of viruses from the Flaviviridae family were identified. Both the right and left sides of the integration were resolved bioinformatically for nrEVEnew-7. Only the left or right sides of the integration were bioinformatically predicted from nrEVEnew-1, nrEVEnew-2 and nrEVEnew-6 or nrEVEnew-4, nrEVEnew-5 and nrEVEnew-8, respectively. nrEVEnew-2, nrEVEnew-4, nrEVEnew-6 and nrEVEnew-7 were detected when running ViR using both AloF1 and AalbF2; nrEVEnew-1, nrEVEnew-5 and nrEVEnew-8 were detected only when running ViR with AalbF2, possibly because of the higher completeness of AalbF2 (28) vs AaloF1 (27). Viral integrations identified running ViR in the genome of *Ae. albopictus* mosquitoes are described in Table 3.

**Table 3.** Viral integrations identified running ViR with WGS data from *Ae. albopictus* mosquitoes.

| nrEVE | Length (bp) | Virus (blastx id %) | Viral protein |
| --- | --- | --- | --- |
| nrEVEnew-1 | 295 | KRV (70%) | NS1-NS2A |
| nrEVEnew-2a | 578 | KRV (82%) | NS3 |
| nrEVEnew-4/6 | 534 | AeFV (75%) | NS5 |
| nrEVEnew-5 | 880 | KRV (78%) | NS5 |
| nrEVEnew-7 | 318 | AeFV (91%) | E |

#### Testing module 2

Running module 1 of ViR led to the identification of both the right and left integration sides of nrEVEnew-7, but without the exact integration points (Fig. 2B). nrEVE-new-7 is a 187 bp sequence with similarity to Aedes Flavivirus (AeFV), with less than 78% nucleotide identity with EVEs already annotated in either AaloF1 (23) or AalbF2 (28). Thus, we implemented ViR-LTFinder to verify whether our WGS data contained reads encompassing the integrations site. Running ViR_LTFinder with our WGS data using as anchor the viral portion of nrEVEnew-7 built to a consensus sequence of 1237 bp including both integration sites (Fig. 2B, section a). ViR_LTFinder was further tested for nrEVEnew-1, nrEVEnew-2, nrEVEnew-4, nrEVEnew-5 and nrEVEnew-6. Results of ViR_LTFinder showed that nrEVEnew-4 and nrEVEnew-6 correspond respectively to the right and left regions of the same integration, for a total of 534 viral bp (Fig. 2B, section b). With ViR_LTFinder the viral sequence of nrEVEnew-2 (Supplementary Fig. 1, step a) was extended for 619 bp to the left predicting an integration break point and 636 bp to the right. The sequence on the right was still viral indicating nrEVE-new-2 is longer (Supplementary Fig. 1, step b). Thus, we repeated ViR_LTFinder as before. This operation produced two consensus sequences: the first one of 1474 bp including both the left and the right anchor sites on the mosquito genome and supporting an integration of 510 bp; the second of 1987 bp extending the sequence obtained in the first step of 577 bp and supporting a longer integration (Supplementary Fig. 1, step c).

#### Molecular validation

PCR-based molecular validation was obtained for all predicted viral integrations, with the exception of nrEVE-new8 and the left integration sites of nrEVE-new-2 (Fig. 2C). We cannot exclude this result is due to the rarity of these integrations. nrEVE-new8 was bioinformatically identified only in one Pool-60 sample. Unfortunately, we were unable to perform PCR on this pool because of the lack of DNA aliquots.

### Testing ViR performance

We tested the benefit of running ViR by calculating the solve dispersion gain parameter. For each sample, we considered as dataset the whole list of reads resulting from ViR_RefineCandidates. We quantified the gain comparing the host-mapping loci identified by VyPER and the read groups created by ViR_SolveDispersion. As an example, in replicate 19 among the single samples, seven pairs of reads were identified by Vy-PER as candidate viral integrations. ViR solved the dispersion of these seven reads by grouping them into one group supporting nrEVEnew-4; two read remained ungrouped, resulting in a Solve Dispersion Gain value of 0,65 (Fig. 3A). In single samples, the median values of the solve dispersion gain were 0.47 and 0.5 when using AaloF1 and Aalb2, respectively. In pool samples, the median values in pool60 were 0.36 and 0.42 and in pool30 were 0.58 and 0.48 when using AaloF1 and Aalb2, respectively (Fig. 3B). Thus, we noticed a significant gain in using ViR, also in the improved version of the *Ae. albopictus* assembly, AalbF2 (28). Dispersion gain values were not different between single vs pools or between pools 30 vs pools 60 samples, indicating the gain is influenced by the sequencing strategy nor the depth of sequencing.

**Figure 3:**
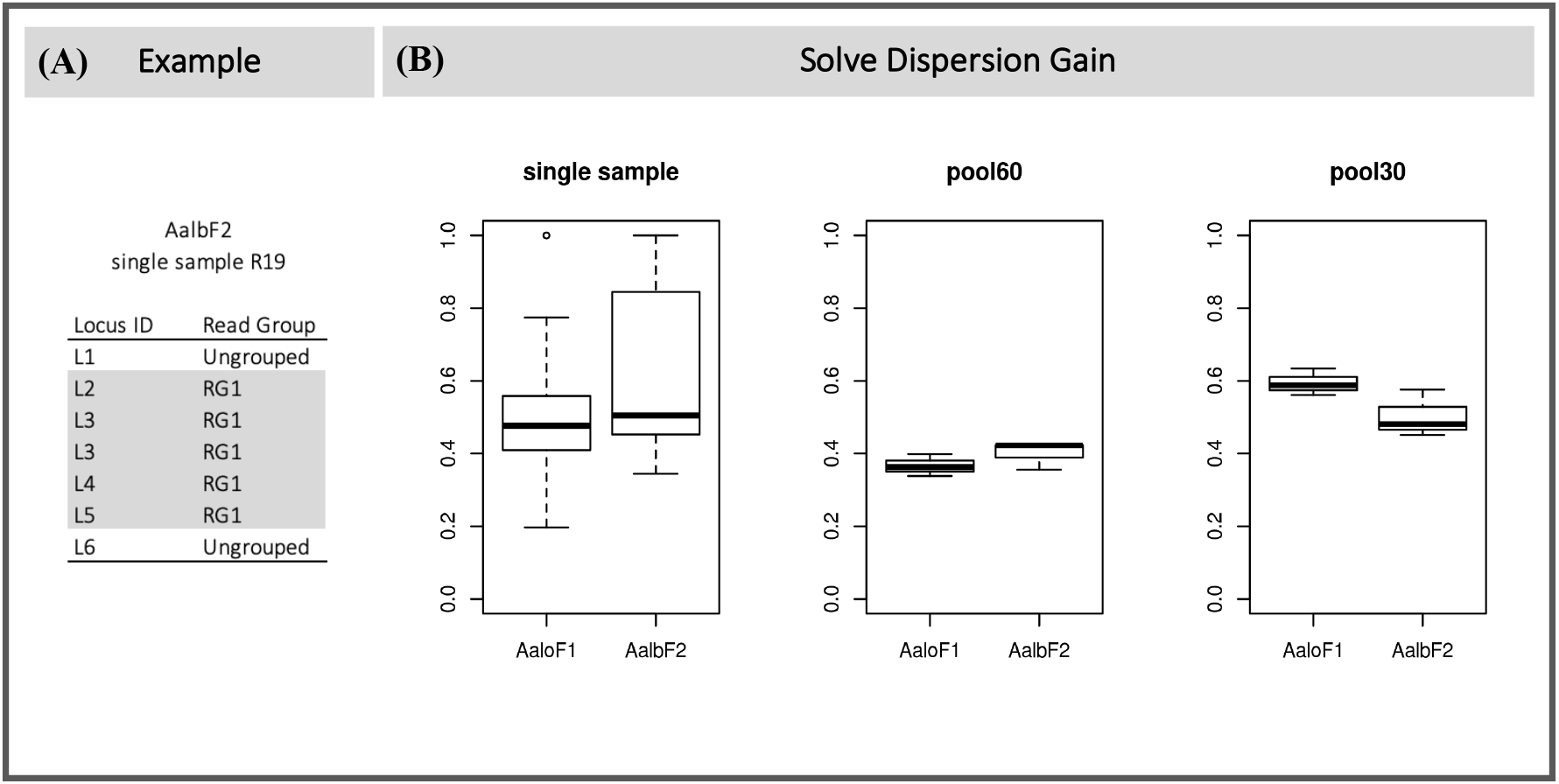
Testing the gain of running ViR. **(A)** Seven pairs of reads were identified by Vy-PER as candidate viral integrations. ViR grouped five of these reads into a single group which led to the identification of nrEVEnew-4 and resulting in a Solve Dispersion Gain value of 0,65. **(B)** Values of Solve Dispersion Gain were calculated for all single and pool samples.

In each dataset a certain number of chimeric reads identified by Vy-PER could not be grouped. These reads have alignments in the genome distant from the others and cannot be used to predict candidate viral integrations. Even if they are not useful for the discovery of viral integrations, it is important to isolate them to avoid wasting time trying to interpret them, for this motif we included these ungrouped reads in the calculation of the Solve Dispersion Gain. These ungrouped reads were more abundant when testing ViR with AaloF1 than AaloF2 as a result of the higher fragmentation of the AaloF1 assembly (28).

## DISCUSSION

The purpose of ViR is to provide reliable identification of EVEs when viral integrations occur in repetitive regions of host genomes or when using genome assemblies of non-model organisms which are still fragmented. Both these elements generate intra-host variability resulting in mis-alignments when trying to map short paired end reads. This issue is magnified in chimeric reads, which are the source of a candidate viral integration because in chimeric reads only one of the pairs maps to the host genome. ViR is not a new pipeline for the identification of EVEs, rather it works to ameliorate predictions. This is important because despite the large number of pipelines currently available to detect EVEs (for a review see (10)), all have been designed for well-annotated model genomes, *in primis* the human genome. The advent of next-generation sequencing technologies and metagenomic analyses facilitated the study of non-model organisms, the genomes of many of which were sequenced showing that LT from viruses may be as frequent in eukaryotes as in prokaryotes (17, 20). Additionally, integrated viral sequences in both plants and animals derive not only from DNA viruses and retroviruses but also from non-retroviral RNA viruses and are enriched in repetitive regions of the genome (20, 23–25, 28). To understand the widespread distribution of EVEs and their biological role, we need to characterize EVEs using WGS data from wild-collected samples or samples collected under hypothesis-driven experimental conditions. In the case of non-model organism, this pursuit requires being able to handle genomes rich in repetitive elements and/or fragmentation in the assembly. Towards this goal, we have developed ViR, a bioinformatic pipeline suited to account for intrasample variability in predicting candidate viral integrations. We show ViR performance by using WGS data from *Ae. albopictus*, the mosquito with the largest genome to date, a TE content >50% of the genome and two assemblies differing in completeness (27, 28). In the absence of a true set of viral integrations, ViR performance was tested calculating the dispersion gain, a parameter from the Information Theory (35) and molecularly validating predicted viral integrations. We demonstrated that ViR is able to solve the dispersion of reads supporting a viral integration similarly when using WGS data from single and pool samples and with a coverage of between 30 and 60x. We anticipate ViR will open new venues to explore the biology of EVEs, especially in non-model organisms. The design of ViR makes it compatible with any list of putative chimeric reads produced by any currently available EVE identification tool, giving users great flexibility. Importantly, while we generated ViR with the identification of EVEs in mind, its application can be extended to detect any LT event providing an *ad-hoc* sequence to interrogate.

## AVAILABILITY

ViR is implemented in bash and python2.7 and is freely accessible on Github.

## SUPPLEMENTARY DATA

Supplementary data will be available in the online journal.

---

**Supplementary Table 1.** Structure of the file “Final_ChimericPairs_Info.txt” which is the output of the script ViR _RefineCandidates. The file will have 18 columns.

| Column Name | Column Description |
| --- | --- |
| SAMPLE_ID | name of the sample |
| READ_ID | ID of the pair of reads |
| HR_CHR | Chromosome in which the host read is mapped |
| HR_START | Start position of the host read |
| HR_END | End position of the host read |
| HR_SEQ | Sequence of the host read |
| HR_FLAG | Flag of alignment of the host read |
| HR_NT_BQ20 | N. of nt with Phred base quality >20 in the host read |
| HR_MQ | Phred mapping quality of the host read |
| VR_SEQ | Sequence of the viral read |
| VR_FLAG | Flag of alignment of the viral read |
| VR_NT_BQ20 | N. of nt with Phred base quality >20 in the viral read |
| VIRUS_ID |  |
| VIRUS_START | ID and coordinates of the virus mapping to the viral read |
| VIRUS_END |  |
| VIRAL_SEQ | viral sequence matching to the viral read |
| VIRAL_SEQ_LEN | Length of the viral sequence |
| VR_AlignToRef? | Y/N flag of alignment of the viral read within the host genome |

**Supplementary Table 2.**
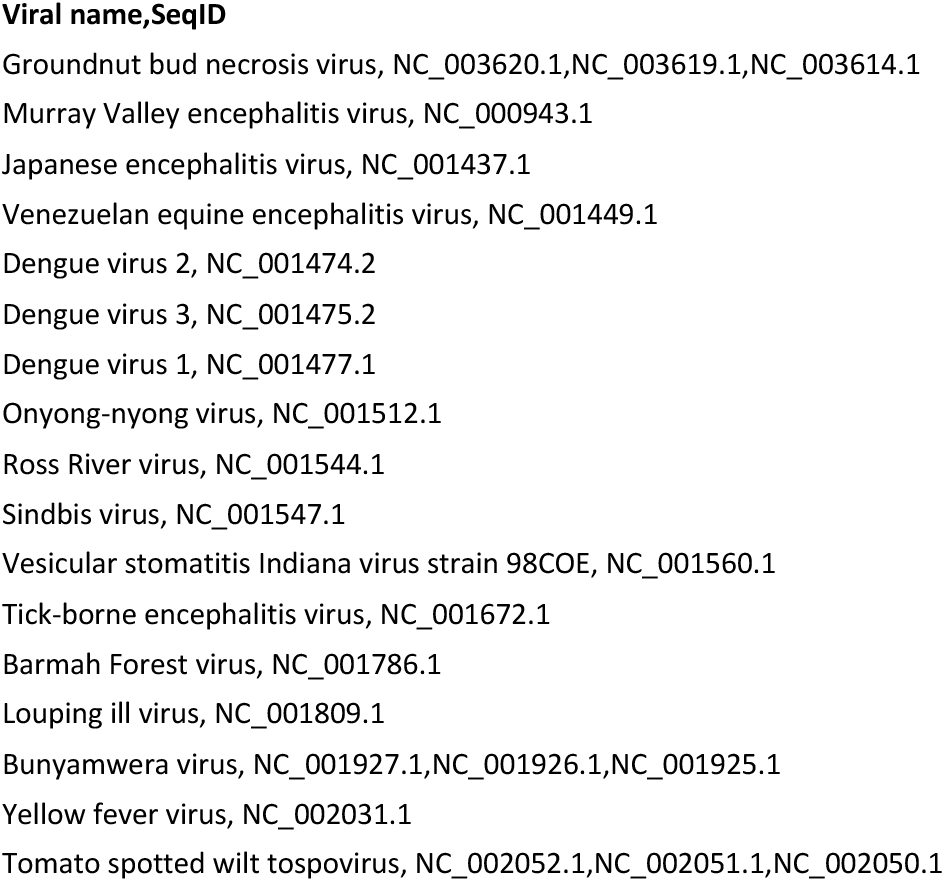
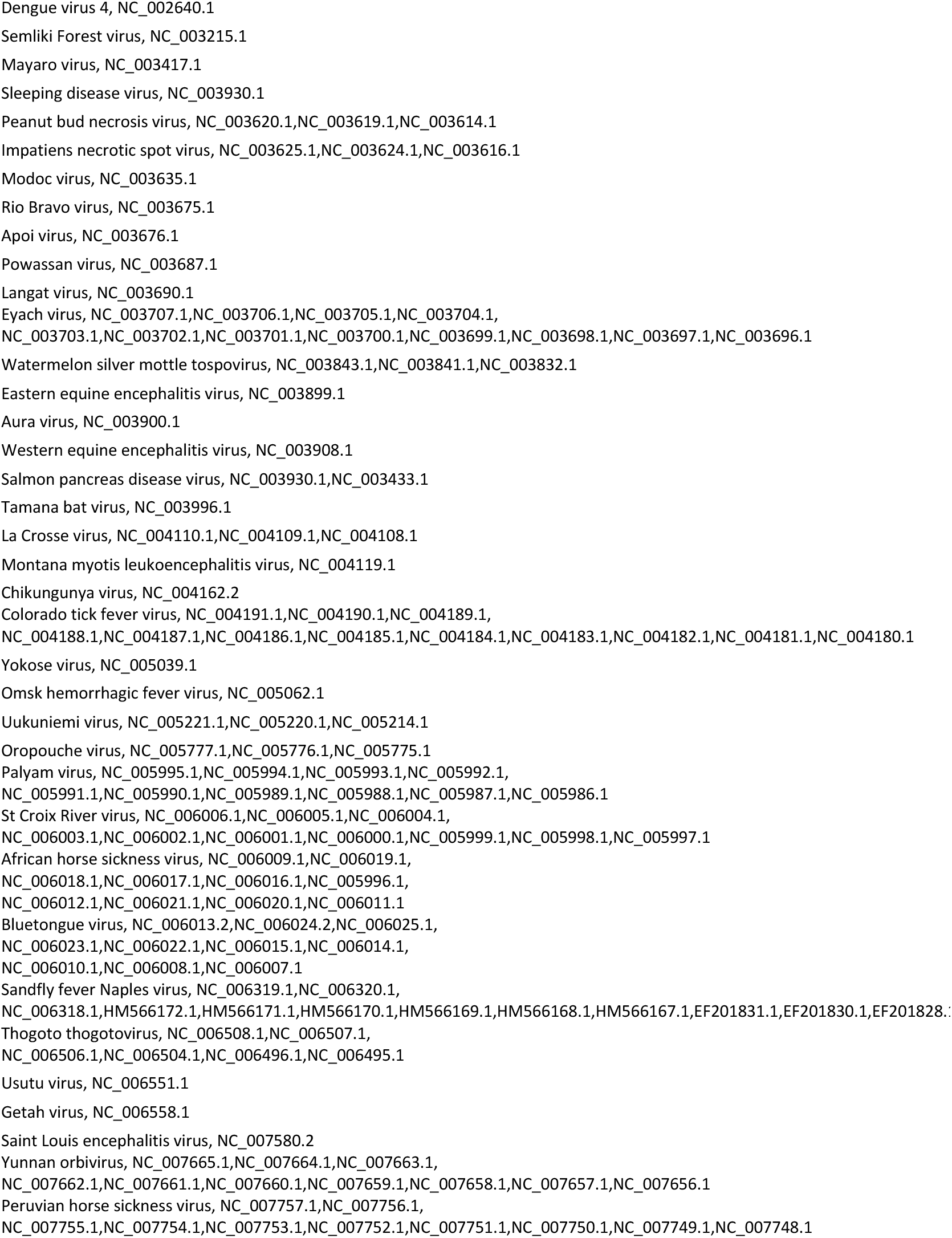
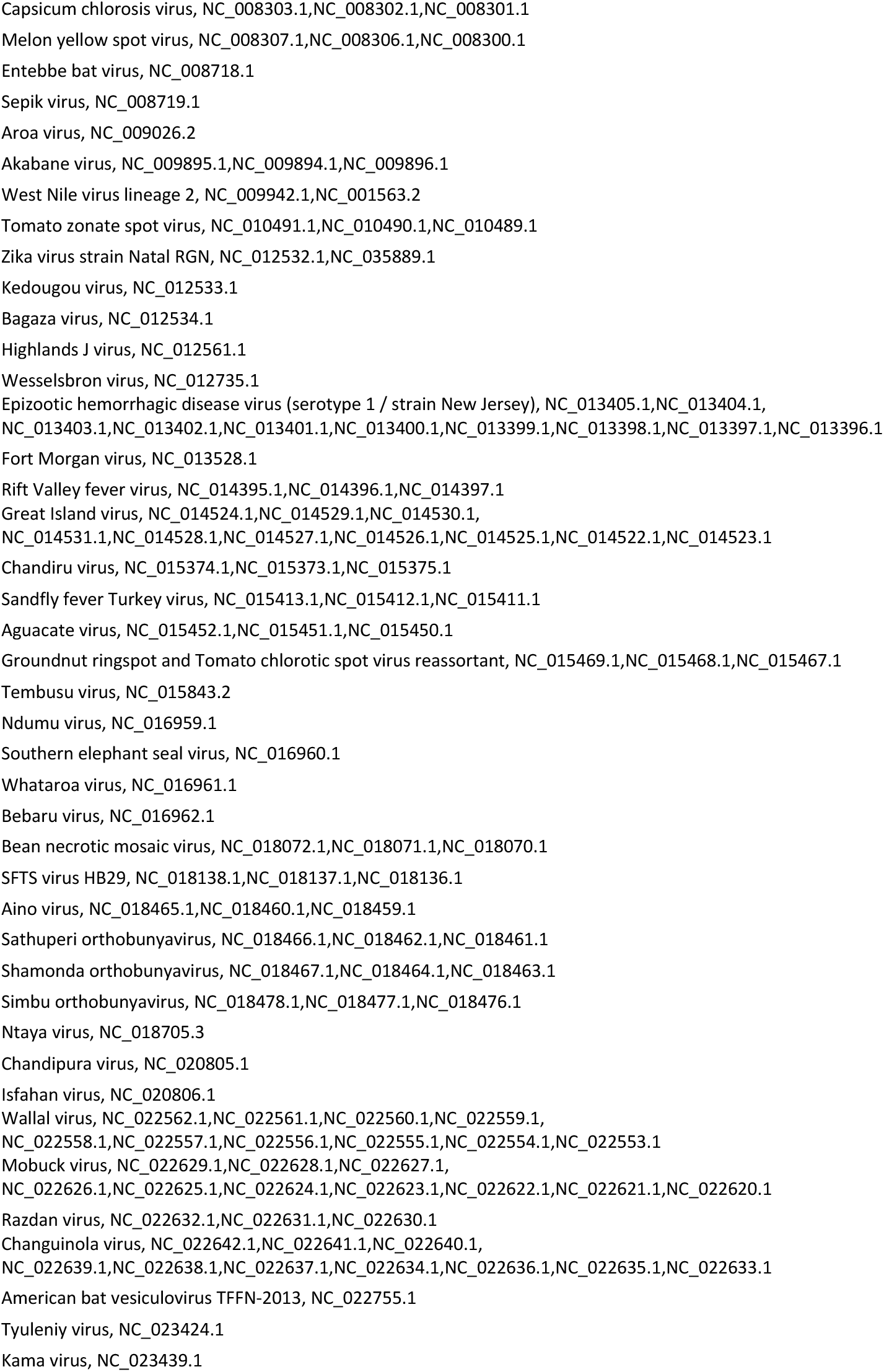
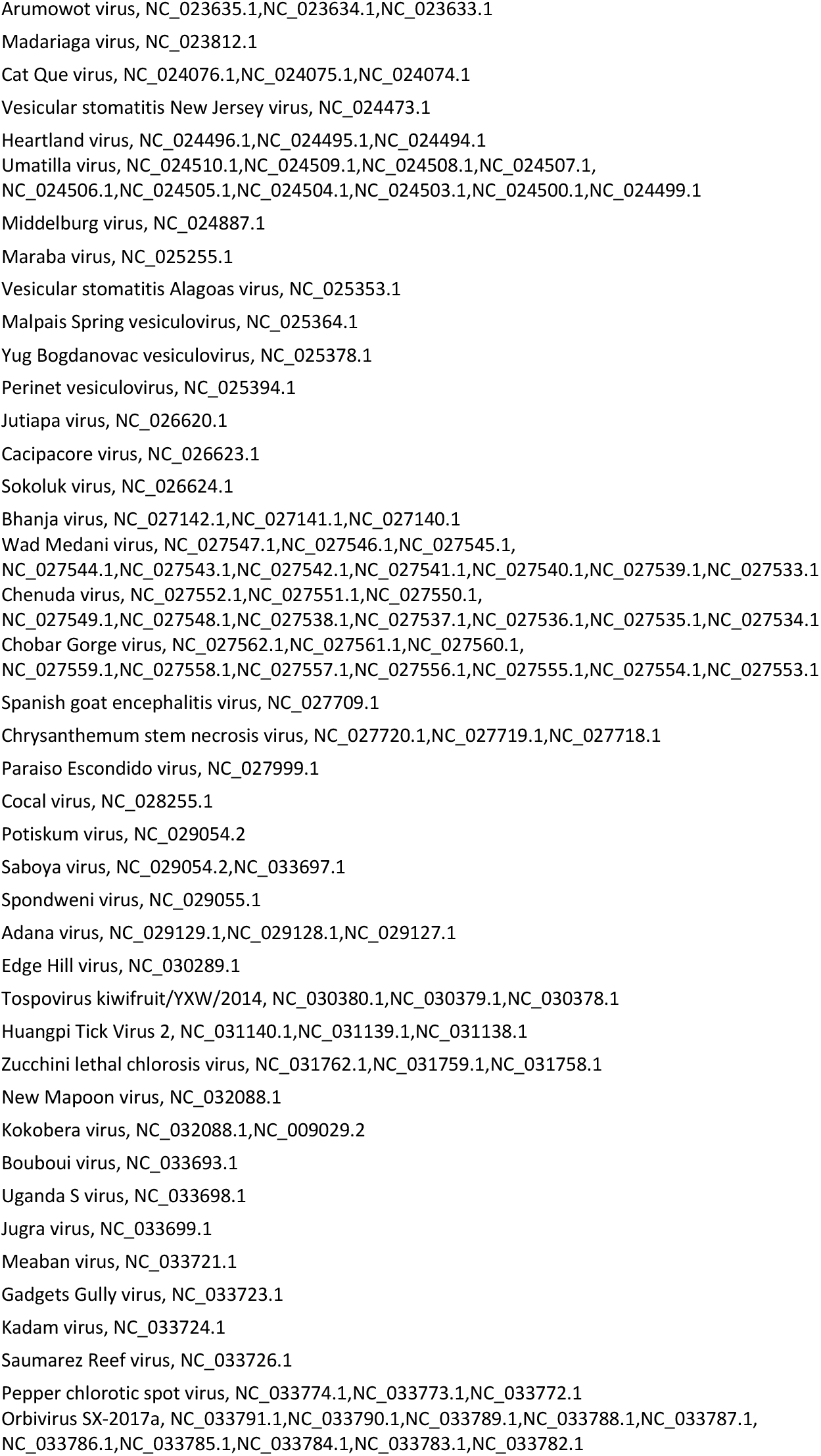
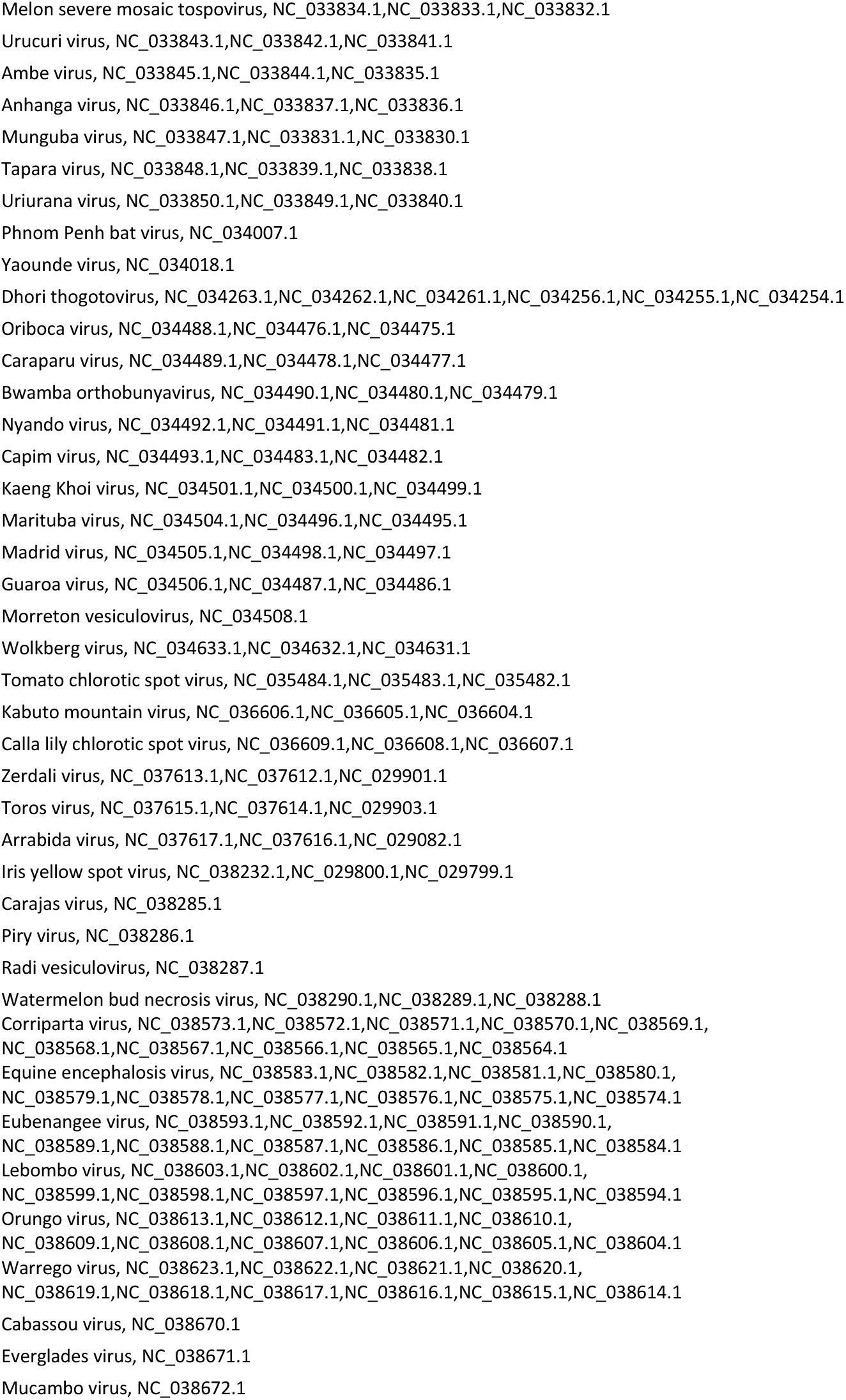
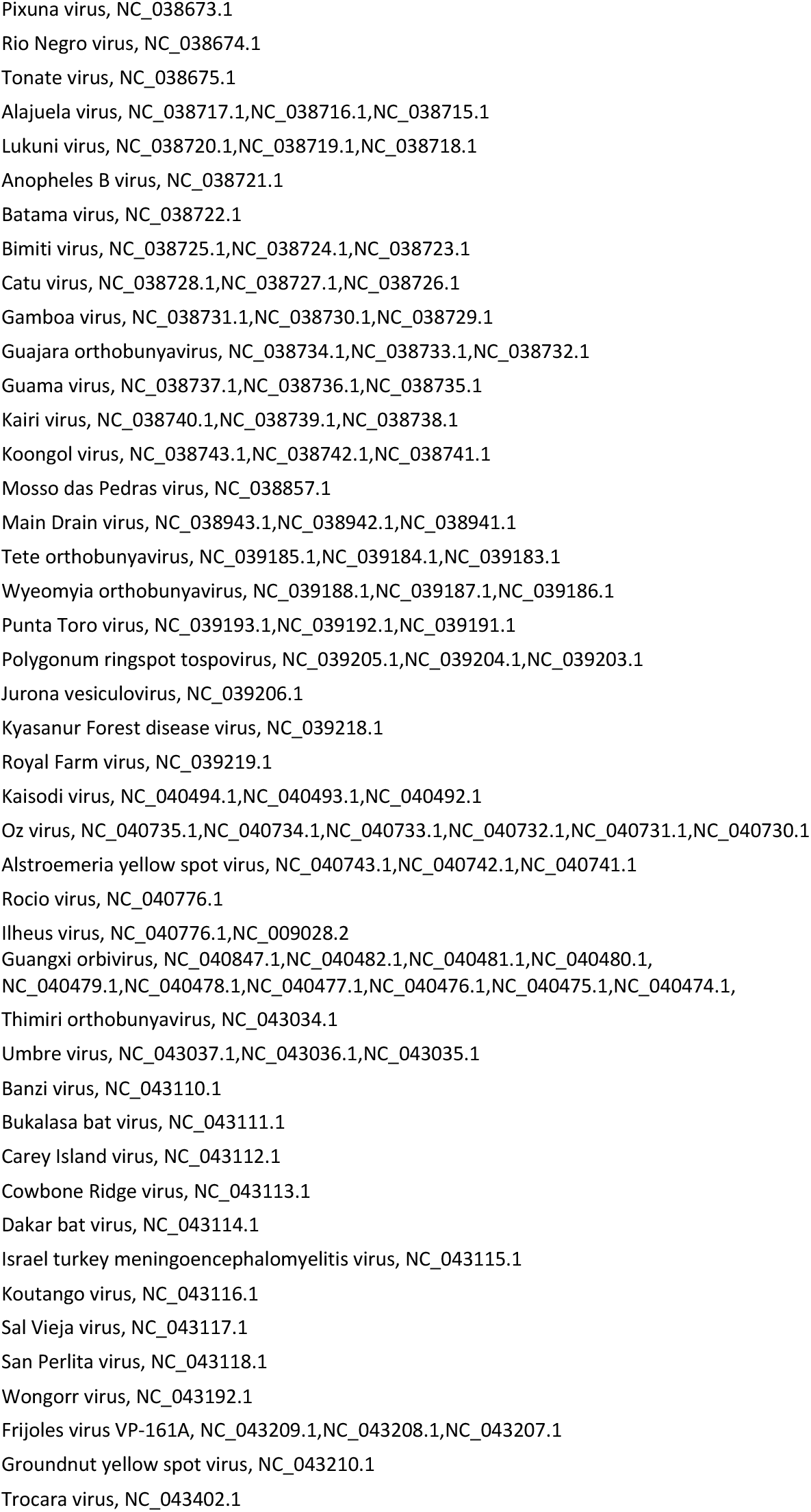
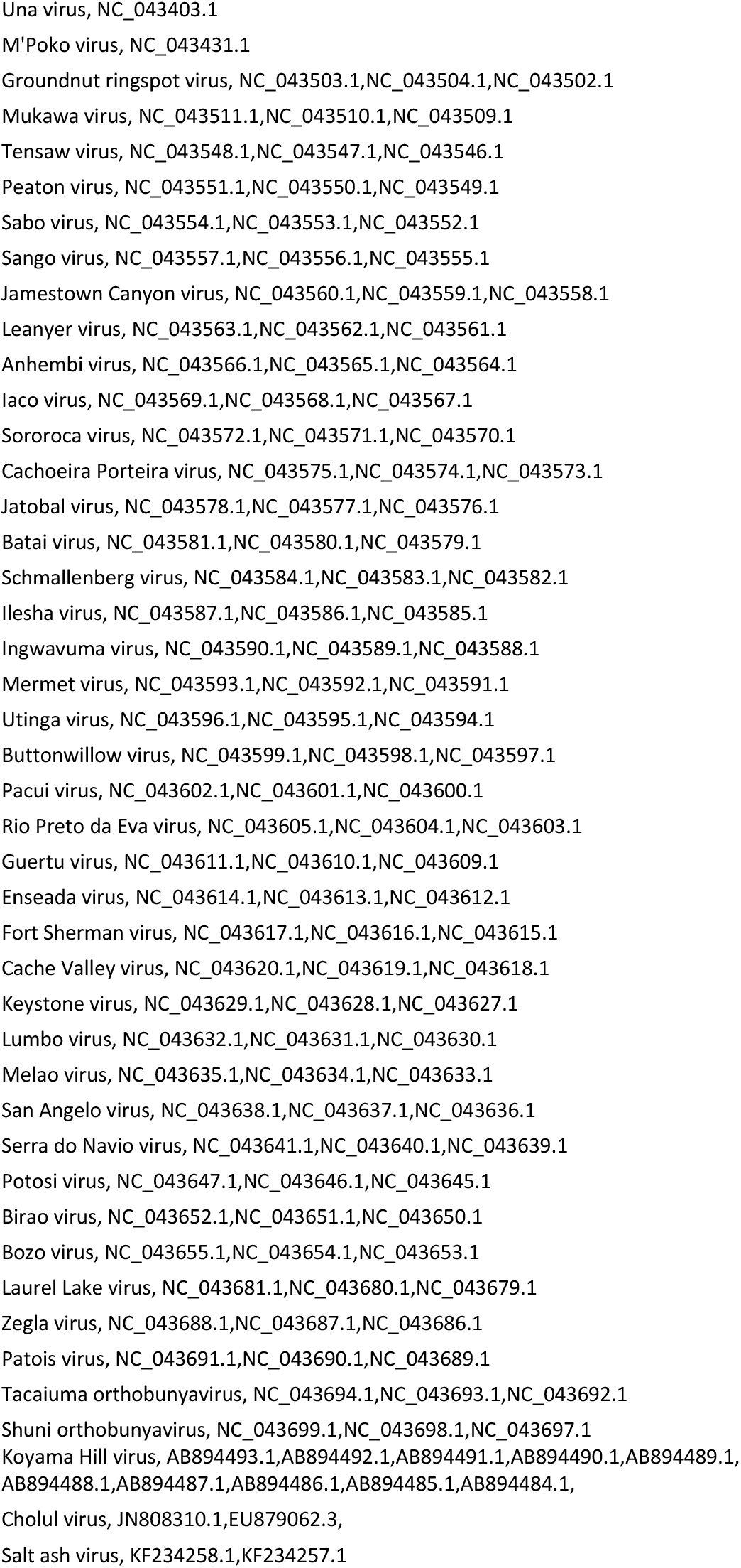
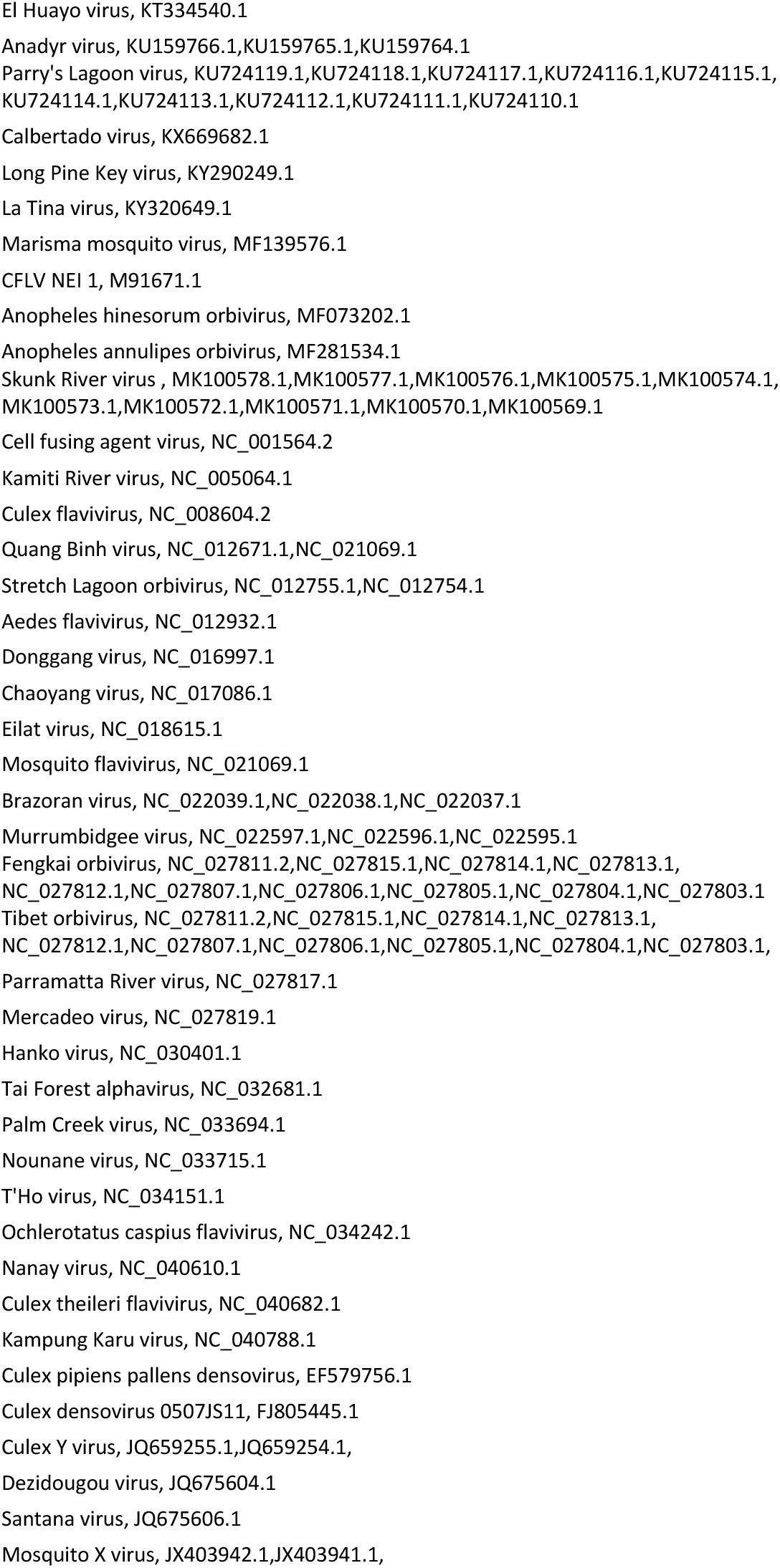
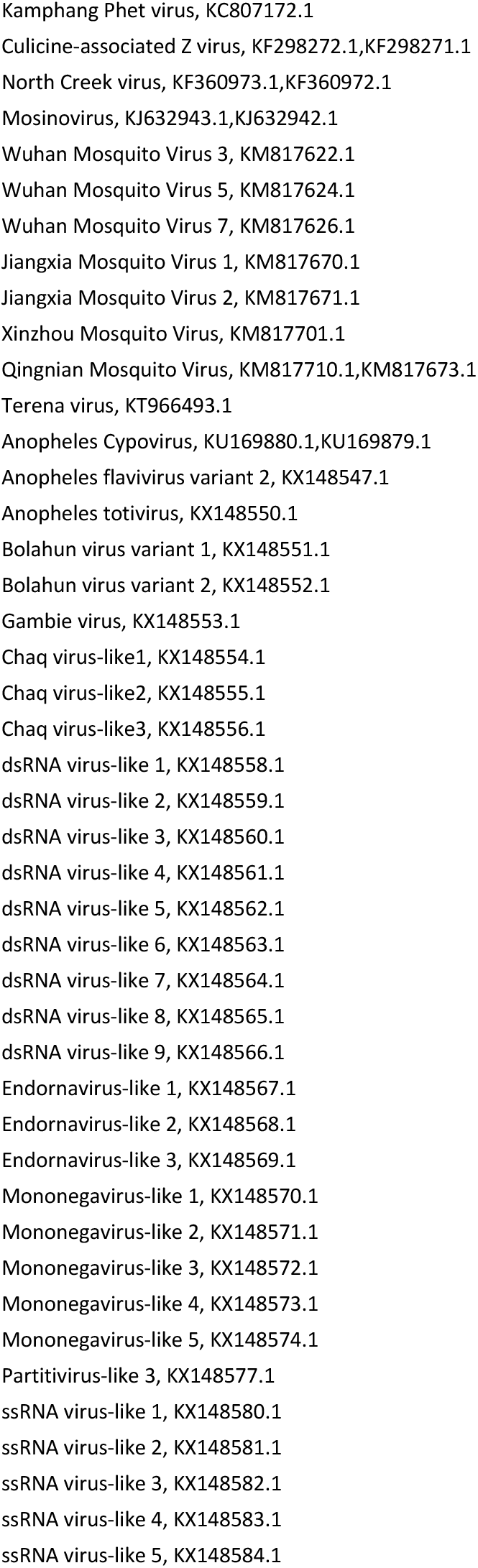
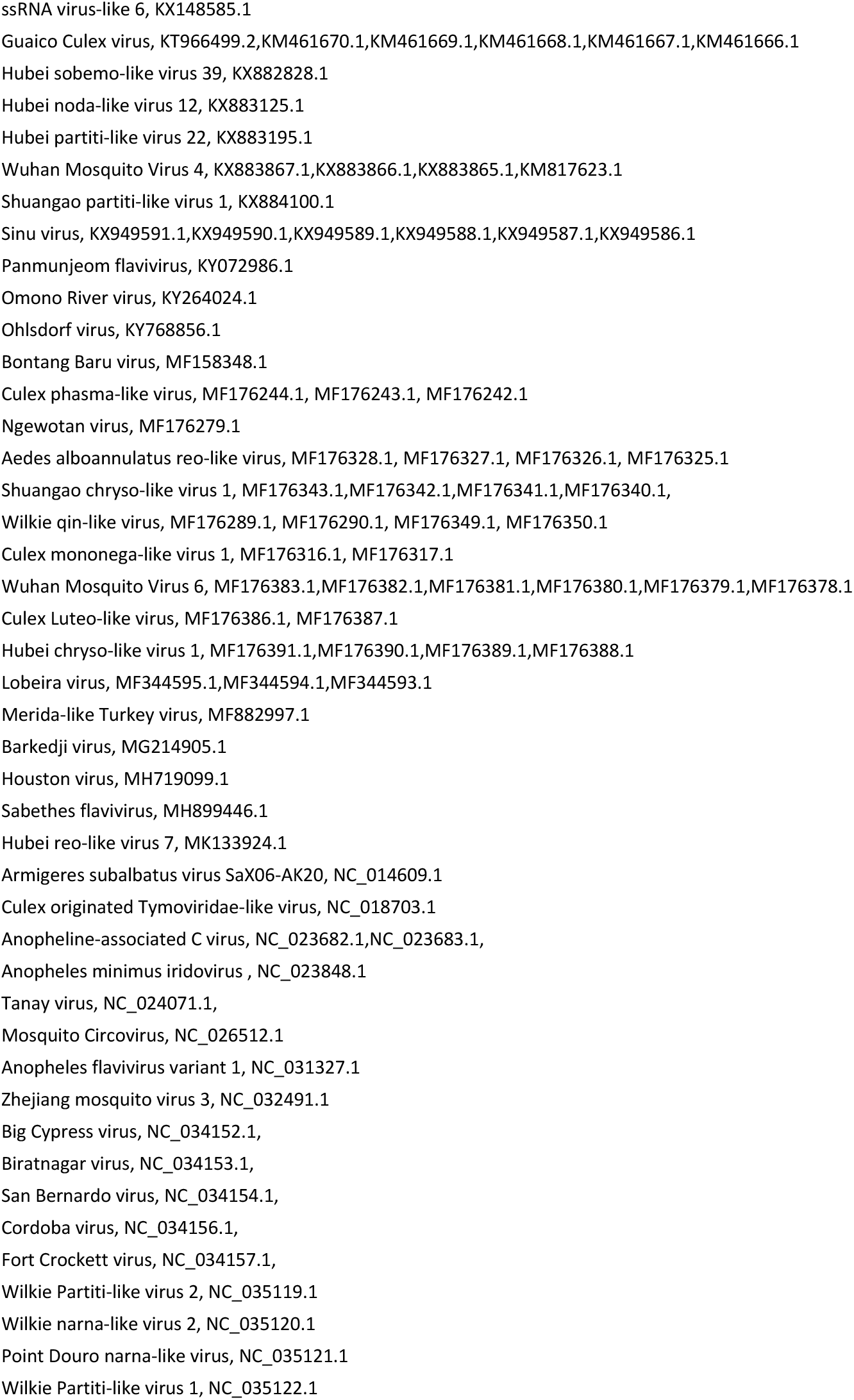
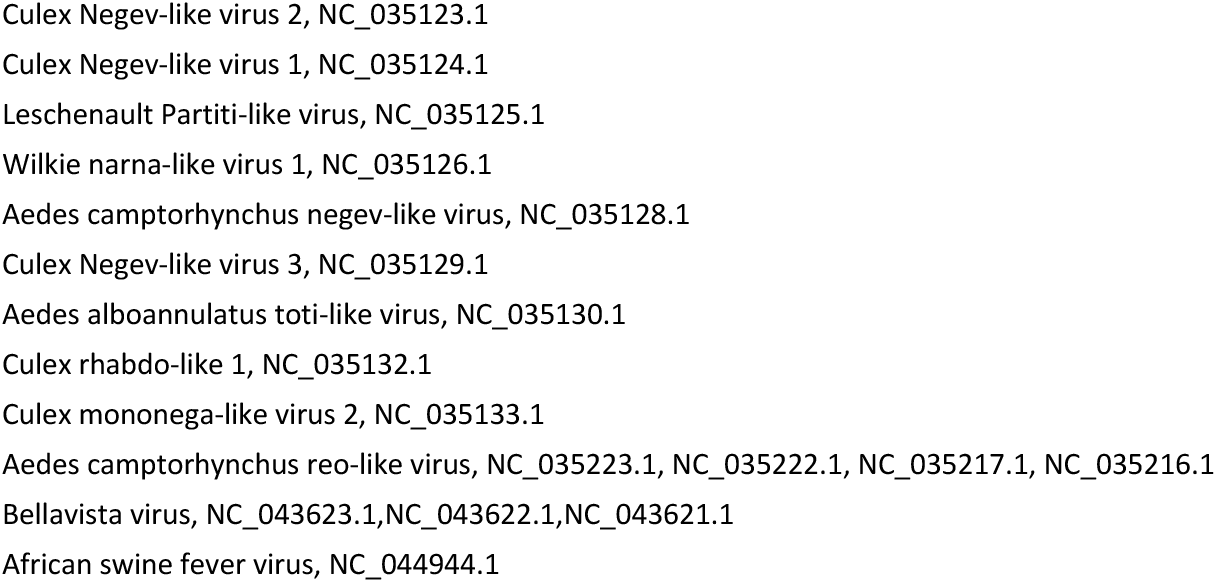
Viral genome database. Viral species and their sequences are shown.

**Sup. Fig. 1.**
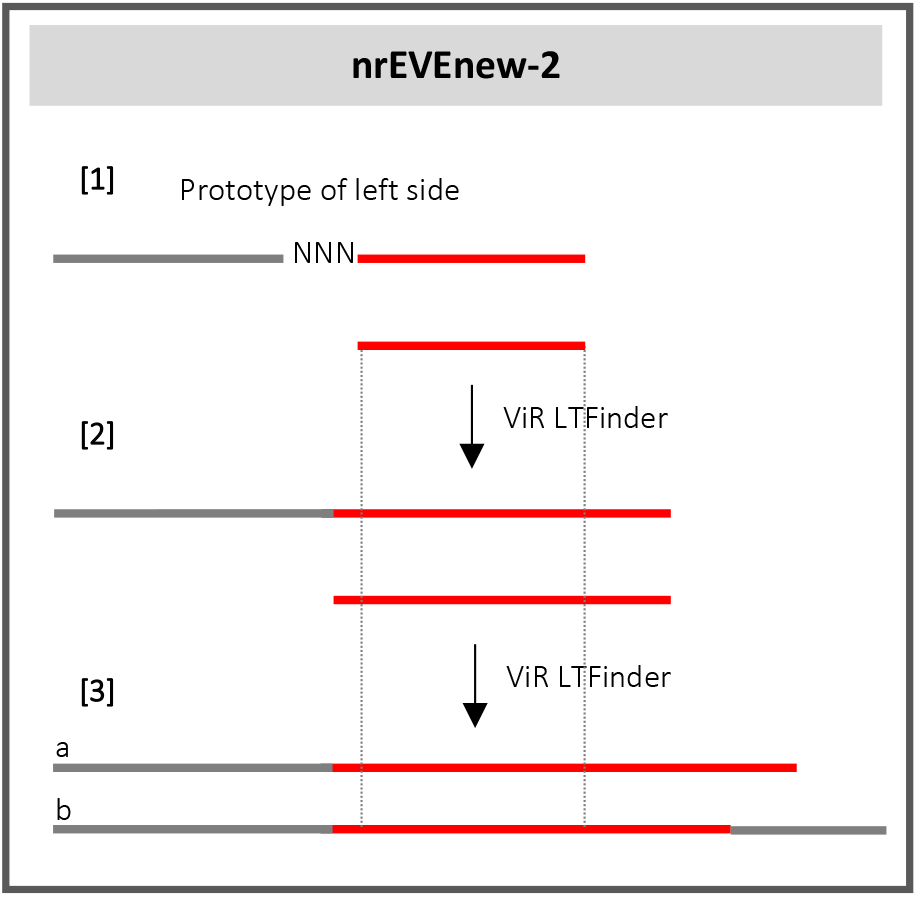

